# The effects of human population density on trophic interactions are contingent upon latitude

**DOI:** 10.1101/2023.08.22.554272

**Authors:** Juan Antonio Hernandez-Aguero, Ildefonso Ruiz-Tapiador, Lucas A. Garibaldi, Mikhail V. Kozlov, Elina Mantyla, Marcos E. Nacif, Norma Salinas, Luis Cayuela

**Affiliations:** Vrije Universiteit Amsterdam; Universidad Politecnica de Madrid; Universidad Nacional de Rio Negro; University of Turku; Pontifical Catholic University of Peru; Universidad Rey Juan Carlos

**Keywords:** bird predation, climate change, global, insect herbivory, latitude, urban heat island effect, urbanization

## Abstract

Aim: Studies conducted at a global scale are necessary to make general conclusions on the effect of urbanization on trophic interactions and explore how these effects change along latitudinal gradients. Since biotic interactions are more intense at lower latitudes, we predict that the intensity of trophic interactions will decrease in response to urbanization (quantified by human population density), with higher impacts of urbanization at higher latitudes.

Location: Global (881 study sites).

Time period: 2000-2021.

Major taxa studied: Birds, arthropods and woody plants.

Methods: We compiled global data on insect herbivory and bird predation from individual studies using similar methodologies, and fitted generalized linear mixed models to test the effect of human population density, latitude and their interaction on these two response variables.

Results: The intensity of herbivory and predation decreased with the increase of human population density at lower latitudes, remained unaffected at intermediate latitudes, and increased at higher latitudes.

Main conclusions: The effect of human population density on insect herbivory and bird predation consistently varies across latitudes, with a reversal of the pattern at high versus low latitudes.

## Introduction

Among human activities that cause habitat loss, urban development produces some of the most significant local extinction rates (McKinney, 2002). At present 54% of the world population live in cities (≥ 300,000 inhabitants) and this percentage is expected to increase up to 70% by 2050 (United Nations, 2018). Cities have recurrent air and water pollution problems and cause profound changes in ecosystems, land use, biogeochemical cycles, and climate (Bai, McPhearson, Cleugh, Nagendra, Tong et al., 2017). Such impacts result in a decline in the abundance and diversity of many taxa, including bats (Russo & Ancilloto, 2015), birds (Gagné & Fahrig, 2011; Planillo, Kramer-Schadt, Buchholz, Gras, von der Lippe et al., 2021), fungi (Abrego, Crosier, Somervuo, Ivanova, Abrahamyan et al., 2020), beetles (Gagné & Fahrig, 2011), bees (Fortel, Henry, Guilbaud, Guirao, Kuhlmann et al., 2014) and other insects (Planillo et al., 2021).

Shifts in species composition, abundance and diversity of biota driven by urbanization can also affect trophic interactions, which are of particular relevance for biological communities and critical ecosystem services, such as nutrient cycling (DeAngelis, 2012), pest control (Whelan, Wenny & Marquis, 2008), adaptation to the effects of climate change or human health (Sirakaya, Cliquet & Harris, 2018). According to the increasing disturbance hypothesis (Eötvös, Magura & Lövei, 2018), urbanization decreases the intensity of trophic interactions, such as herbivory or predation. For herbivory, evidence in support of this hypothesis have been found in Europe (Kozlov, Lanta, Zverev, Rainio, Kunavin et al., 2017; Moreira, Abdala - Roberts, Berny Mier y Teran, Covelo et al., 2019), and USA (Meineke, Classen, Sanders, & Davies, 2019). In addition, a recent review article concludes that urbanization could affect both adaptive and non-adaptive evolution of herbivorous arthropods and their host plants in urban environments, altering plant-herbivore interaction, yet changes in abundance of native species appear to be species specific, without general consensus on their effects (Miles, Breitbart, Wagner & Johnson, 2019). For predation, evidence in support of the ‘increasing disturbance’ hypothesis includes the reduction in abundance of predators in response to urbanization in southern England (Rocha & Fellowes, 2018), or the decrease in predation in Finland (Jokimäki & Huhta, 2000), Denmark (Ferrante, Cacciato & Lövei, 2014), Madagascar (Schwab, Wurz, Grass, Rakotomalala, Osen et al., 2020), North America (Gering & Blair, 1999; Thorington & Bowman, 2003) and Europe (Eötvös, Magura & Lövei, 2018). Despite widespread support for the ‘increasing disturbance’ hypothesis, some studies have also reported a positive effect of urbanization on herbivory in different parts of the world, including Brazil (Cuevas-Reyes, Gilberti, González-Rodríguez, & Fernandes, 2013; Rivkin & Moura, 2020), Australia (Christie & Hochuli, 2005) and USA (Cregg & Dix, 2001; Dale & Frank, 2017). Likewise, predation by birds and arthropods has been shown to increase across disturbance gradients in Philippines (Posa, Sodhi, & Koh, 2007) and in Europe (Kozlov et al., 2017). Different hypotheses might explain these contrasting results. First of all, the ‘predator relaxation/safe habitat’ hypothesis (Noske, 1998) proposes that there are less predators in disturbed areas, which results in less mortality of prey and, consequently, an increase in herbivory. This reduces the need for vigilance behavior, allowing the increase of other activities such as feeding or reproduction. Similarly, the ‘natural enemy release’ hypothesis (Keane & Crawley, 2002) proposes that urban areas, having less complex vegetation, may support fewer natural enemies, decreasing the biological control of pests, which allows herbivore species to proliferate (Dale & Frank, 2017; Meineke et al., 2014). Also, the ‘plant stress’ hypothesis proposes that herbivore species could proliferate in urban areas due to a reduction in defense investment of plants explained by water stress, elevated temperature and pollution (White, 1969). At the same time, the ‘plant vigor’ hypothesis explains the increase in herbivory due to the increase of growth of plants due to the fertilizing effect of nutrients and CO_2_ in urban areas (Price 1991). Alternatively, the ‘predation proliferation’ hypothesis (Sorace, 2002) proposes that certain predators can be adapted to urban environment, the so-called ‘urban exploiters’ in contrast to ‘urban avoiders’ (sensu Blair 1996). This increase of abundance added to the presence of exotic predators (Sasvári et al., 1995) can conduct to an increase in predation pressure.

The above-mentioned studies conducted at regional or local scales illustrate that the effects of urbanization on both herbivory and predation are context-dependent and vary across regions. This variation could be associated with latitudinal gradients in biotic interactions, which at lower latitudes are more intense than at higher latitudes (Zvereva & Kozlov, 2021). This pattern might be explained by the ‘latitudinal biotic interaction’ hypothesis (Dobzhansky, 1950), which proposes that primary productivity and species richness increase in the tropics (Dobzhansky, 1950; Novotny et al., 2006), mostly as a result of warm temperatures all year-round (Coley & Aide, 1991). The consequences of this richness increase are that more productive ecosystems sustain larger levels of herbivory (McNaughton et al., 1989), while more diverse predator communities create redundancies and complementarity in prey consumption, which results in an increase in predation (‘Paine’s predator’ hypothesis; Paine, 1966). Despite the theoretical and empirical background supporting the ‘latitudinal biotic interaction’ hypothesis (e.g. Pennings & Sillman, 2005; Pennings Ho, Salgado, Więski, Davé et al., 2009; Garibaldi, Kitzberger & Ruggiero, 2011; Moreira, Abdala - Roberts, Parra - Tabla, & Mooney, 2015), the literature has contradictory evidence about the effect of latitude on the intensity of biotic interactions (Anstett, Nunes, Baskett & Kotanen, 2016). In the particular case of herbivory, some studies point out to an increase in insect herbivory with latitude (Adams & Zhang, 2009; Del-Val & Armesto, 2010), while others report no latitudinal differences (Salazar & Marquis, 2012) or a peak at intermediate latitudes (Kozlov, Lanta, Zverev & Zvereva, 2015). In predation, Lövei & Ferrante (2017) found no effects of latitude on the strength of biotic interactions, whereas other studies found different and even opposite effects depending on the predator (Roslin, Hardwick, Novotny, Petry, Andrew et al., 2017; Zvereva, Castagneyrol, Cornelissen, Forsman, Hernández-Agu[ero et al., 2019).

Studies conducted at a global scale are necessary to produce general conclusions about the effect of urbanization on trophic interactions and to understand how these effects change along the latitudinal gradients. To date, only a few studies have tested the interaction between latitude and urbanization on herbivory (Kozlov et al., 2017; Meineke et al., 2019; Moreira et al., 2019; Valdés-Correcher, Popova, Galmán, Prinzing, Selikhovkin et al., 2022) and none on predation. These studies conclude that the effects of urbanization on herbivory do not change along the latitudinal gradient. Yet, these studies comprise small latitudinal gradients (10° to 20° latitudinal range) or use non-comparable categorical variables (urban – rural) as a proxy of urbanization, which made their conclusions limited. To fill in this research gap, we investigated the effects of human population density—as a continuous proxy of urbanization—on trophic interactions (insect herbivory and bird predation) along a 122° latitudinal gradient. To achieve this goal, we compiled data on trophic interactions between plants, insects, and birds all over the world, which were collected using similar (and thus comparable) protocols. Using a broad latitudinal gradient encompassing both hemispheres, we attempted to identify whether the effects of human pressure on trophic interactions are universal or change latitudinally. We expected the intensity of trophic interactions to decrease in response to urbanization, as predicted by the ‘increased disturbance hypothesis’, with higher impacts of urbanization at higher latitudes. To our knowledge, this is the first study exploring latitudinal variation in the effects of urbanization on trophic interactions at a global scale.

## Methods

### Herbivory

We extracted most of our herbivory data from a systematic global review (Kozlov et al., 2015), and we searched for additional data published between 2015 and December 2021 in the ISI Web of Science on 19 April 2022 using as keywords “Defoliation” and “Insect herbivory [i.e. Defoliation (All Fields) AND insect herbivory (All Fields) and 2021 OR 2020 OR 2019 OR 2018 OR 2017 OR 2016 OR 2015 (Publication Years)]”. "Defoliation” was used to avoid including other types of herbivory such as galler herbivory or miner herbivory, while “insect herbivory” was included to avoid papers referring to vertebrate herbivores. Since human population density is changing rapidly, we restricted our analysis to data collected after 2000. We selected all papers where insect herbivory was measured as the proportion of leaf area consumed by defoliating insects, either through image software or visually, as proposed by Alliende (1989). Galler and miner herbivory were excluded from our analyses because they were not systematically included in the reviewed papers (e.g. Mendes et al., 2021). We only included data of herbivory on woody plants following the same searching criteria as Kozlov et al. (2015). From the 131 articles obtained only eight used the selected methodology to quantify herbivory. We also included the data provided in Hernández-Agüero, Ruiz-Tapiador, Garibaldi, Kozlov, Nacif et al. (2023) (Figure 1a). A list of the data sources (10 sources) can be found in Supplementary Material 1 Table S1, and a complete list of the studies reviewed (130 sources) and the criteria used for exclusion can be found in Supplementary Material 2.

**Figure 1.**
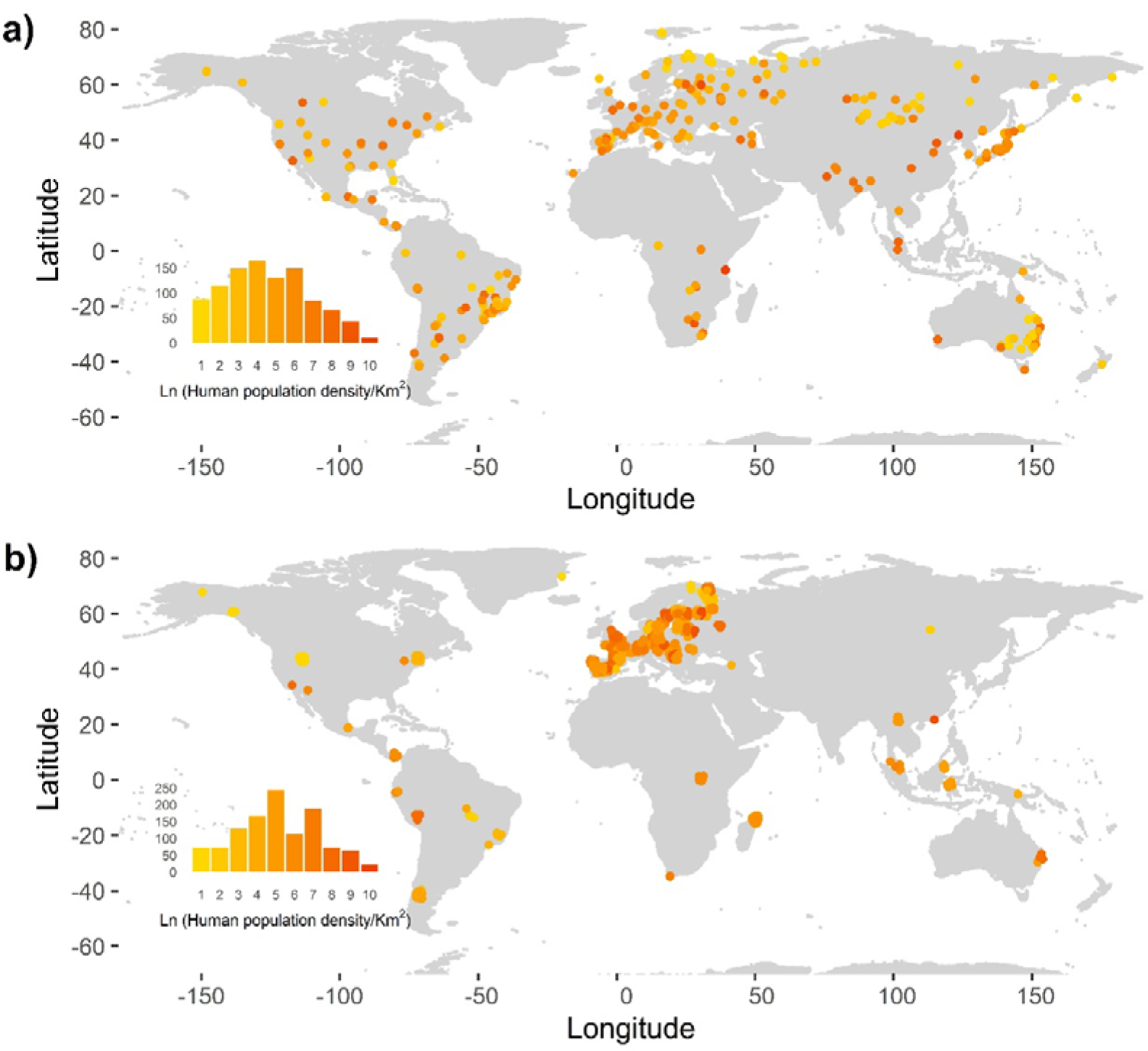
Location of data from selected studies on a) herbivory (n = 326 localities) and b) predation (n = 374 localities). The yellow-red scale represents the natural logarithm of human population density. The histogram located at the bottom left of each figure represents the frequency of human population density on a ln-scale.

### Predation

We extracted predation data from two global studies (Roslin et al., 2017; Zvereva et al., 2019) and searched for additional studies (published between January 2000 and 23 December of 2021) in the ISI Web of Science on 19 April 2022, using the keywords “bird”, “predation” and “larvae” or “caterpillar”[i.e. Results for ‘bird (All Fields)’ AND ‘predation (All Fields)’ AND ‘larvae (All Fields) OR caterpillar (All Fields)’ and ‘2021 OR 2020 OR 2019 OR 2018 OR 2017 OR 2016 OR 2015 OR 2014 OR 2013 OR 2012 OR 2011 OR 2010 OR 2009 OR 2008 OR 2007 OR 2006 OR 2005 OR 2004 OR 2003 OR 2002 OR 2001 OR 2000 (Publication Years)’]. Bird predation on herbivorous insects was quantified by attack rates on artificial caterpillars made of odorless plasticine, which were placed on a woody plant branch or leaves. Following an exposure (from 2 to 64 days), the caterpillars were revisited and bird marks were counted following the methodology proposed in Low, Sam, McArthur, Posa & Hochuli (2014). Most of the studies identified included caterpillars of different colors, except for Roslin et al. (2017) which was based only on green-colored caterpillars. Because prey colors can influence attack rates (Hernández-Agüero, Polo, García, Simón, Ruiz-Tapiador et al., 2020) and colour preferences by birds can change latitudinally, we selected only green caterpillar data, which does not show changes in preference by predators along latitudinal gradients (Zvereva et al., 2019). Our search identified 217 publications, among which only 11 studies shared similar methods to those of Roslin et al. (2017) and Zvereva et al. (2019) and provide raw data on green plasticine models; we also included the data from the submitted manuscript by Hernández-Agüero et al. (2023) (Figure 1b). A list of the data sources (16 sources) is found in Supplementary Material 1 Table S2, and a complete list of the studies reviewed (320 sources) and the criteria used for exclusion can be found in Supplementary Material 2.

To account for differences across studies in the length of the study period, we estimated the probability of a caterpillar to be attacked over one day (henceforth probability of bird predation) as:

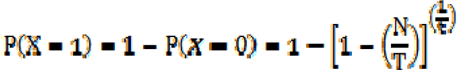

where P(X=1) is the probability of having one caterpillar attacked by birds, *N* is the number of attacked caterpillars, *T* is the total number of caterpillars used per period, and *t* is the period length in days.

### Human population and latitude

We used the Gridded Population of the World (GPWv4 2020) dataset, at 1 km spatial resolution, to estimate human population density at each study site, where each site contained one to several observations of either herbivory or predation. To do so, we created a 10 km radius buffer at each site with the ‘st_buffer’ function of the ‘sf’ package (Pebesma, 2018). This buffer size has been used in other studies to investigate the effects of human impact on vertebrate and plant species (Kim, Mizuno & Kobayashi, 2003; Pautasso, 2007) and is highly correlated (r ≥ 0.7) with values of population density obtained at smaller (e.g. 5 km radius buffer) and larger (e.g. 25 or 40 km radius buffer) scales (Supplementary Material 1 Figure S1 and Figure S2). We then cropped the rasterized GPWv4 dataset with our site buffer of 10 km using the ‘crop’ function of the R package ‘raster’ (Hijmans, 2020) and created a new raster object using our buffered coordinates and the GPWv4 cropped dataset with the ‘mask’ function. Finally, we extracted the total human population at every site with the ‘extract’ function and then human population density per km^2^ was calculated by dividing by the area in km^2^. Human population density was correlated with other commonly used indices of urbanization, such as the proportion of built area (Pearson’s r = 0.73, n = 915, p < 0.001; Supplementary material 1 Figure S3), which was obtained from the Dynamic World V1 (Brown et al., 2022). We used absolute (i.e. unsigned) latitude as a subrogate of climatic conditions, which was highly correlated with mean annual temperature (Pearson’s r = −0.83, n = 915, p < 0.001; Supplementary material 1 Figure S3). Mean annual temperature was estimated at each study site as the averaged values between 1970 and 2000. Climatic data were obtained from the WorldClim database (Fick & Hijmans, 2007) using the function ‘getData’ from the ‘raster’ package. No correlation was found between human population density and absolute latitude (Pearson’s r = −0.07, n = 915, p = 0.023; Supplementary material 1 Figure S3). Results of correlations for herbivory (Figure S4) and predation (Figure S5) can be found in Supplementary material 1.

### Data analyses

We used generalized linear mixed models (GLMM) with a beta error distribution and a logit link function to investigate the effects of human population and latitude on bird predation and insect herbivory. Beta is a family of continuous probability distributions defined on the interval [0,1], and therefore appropriate for the type of response variables we are modelling in this study. We used the natural logarithm of human population, absolute latitude, and their interaction as predictors. In the GLMMs for herbivory, we included the following random factors: i) site, which accounted for potential spatial autocorrelation (there were from 1 to 49 data per site); and ii) plant species nested within genus, which accounted for differences in palatability and plant defenses against herbivory among plant taxa. In the GLMMs for bird predation, we included site as a random factor (there were from 1 to 12 data per location). All GLMMs were fitted using maximum likelihood with the function ‘mixed_model’ of the R package ‘glmmTMB’ (Brooks, Kristensen, van Benthem, Magnusson, Berg et al., 2017).

For both herbivory and predation, alternative models were compared using the Akaike information criterion (AIC) to evaluate the effects of explanatory variables (i.e. fixed effects). Models with a difference in AIC > 2 indicated that the worst model could be omitted. Following Nakagawa & Schielzeth (2013), we estimated the R^2^ of all plausible linear or mixed models. This allowed two components of R^2^ to be calculated: (1) a marginal R^2^ (R^2^_m_) that only considers the variability explained by fixed effects; and (2) a conditional R^2^ (R^2^_c_) that accounts for the variability supported by both fixed and random effects. Model residuals were explored using a simulation-based approach to create readily interpretable scaled (quantile) residuals for the fitted GLMMs (Hartig, 2019). Moran’s index was used to estimate spatial autocorrelation in model residuals both for proportion of herbivory and probability of predation. To test for significance of spatial autocorrelation, this index was compared with a null model random distribution using the ‘spdep’ package (Bivand & Wong, 2018). When spatial autocorrelation in model residuals was significant, we re-fitted the model including a spatial autocorrelation function with exponential correlation structure. This uses an Euclidean distance matrix based on site coordinates (Brooks et al., 2017). We repeated all the analyses with human population density calculated at different buffer sizes (i.e. 1, 5, 10, 15, 20, 25, 30, 35 and 40 km radius buffer) to ensure that the spatial scale at which human population density was measured did not affect the outcome of our analyses (Supplementary Material 1 Table S3 and Table S4).

## Results

### Herbivory

We combined information on herbivory from 508 woody plant species of 271 genera and 113 families collected in 408 different localities spanning 122° latitude (from 43° S to 79° N), with human population ranging from 0 to 21,899 inhabitants/km^2^. The mean proportion of leaf area consumed was 0.0705 (CI_95_: 0.0644 - 0.0765). The best model to explain the variability observed in herbivory included human population density on ln-scale (10 km radius buffer), absolute latitude and their interaction (Table 1; Supplementary material 1 Table S5). Values of conditional R^2^ (0.763) compared to marginal R^2^ (0.046) shows the strong effect of random effects (plant species) to determine herbivory pressure. Model residuals did not show spatial autocorrelation (Moran I statistic standard deviate = −1.6568, p = 0.9512).

**Table 1:**
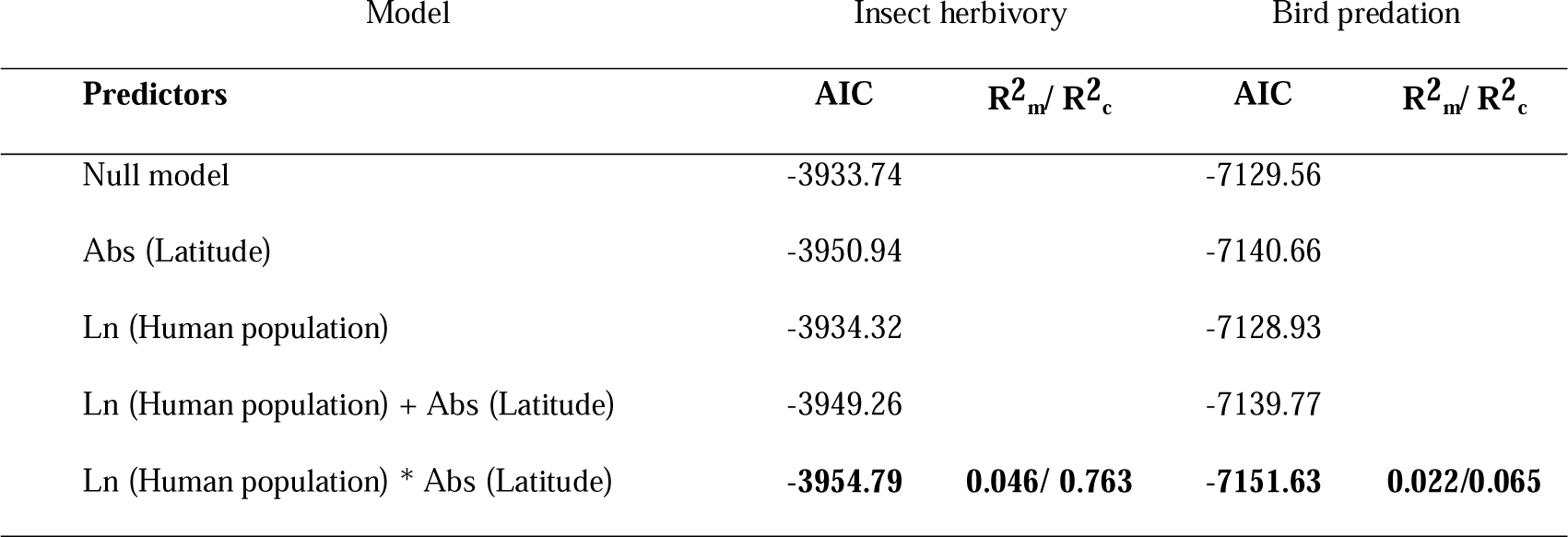
Comparison of alternative models using the Akaike information criterion (AIC). The best model (lowest AIC) is indicated in boldface type. R^2^_m_: a marginal R^2^ that only considers the variability explained by fixed effects; R^2^_c_: conditional R^2^ that accounts for the variability supported by both fixed and random effects.

| Model | Insect herbivory |  | Bird predation |  |
| --- | --- | --- | --- | --- |
| Predictors | AIC | $R^2_m / R^2_c$ | AIC | $R^2_m / R^2_c$ |
| Null model | -3933.74 |  | -7129.56 |  |
| Abs (Latitude) | -3950.94 |  | -7140.66 |  |
| Ln (Human population) | -3934.32 |  | -7128.93 |  |
| Ln (Human population) + Abs (Latitude) | -3949.26 |  | -7139.77 |  |
| Ln (Human population) * Abs (Latitude) | **-3954.79** | **0.046/ 0.763** | **-7151.63** | **0.022/0.065** |

Model predictions showed differential effects of human population density on insect herbivory along the latitudinal gradient. In lower latitude localities (< 25°), herbivory decreased as human population increased. This effect disappeared towards more temperate latitudes (∼ 35°) and reversed at higher latitudes (> 45 °), i.e. higher herbivory in more populated areas (Figure 2).

**Figure 2:**
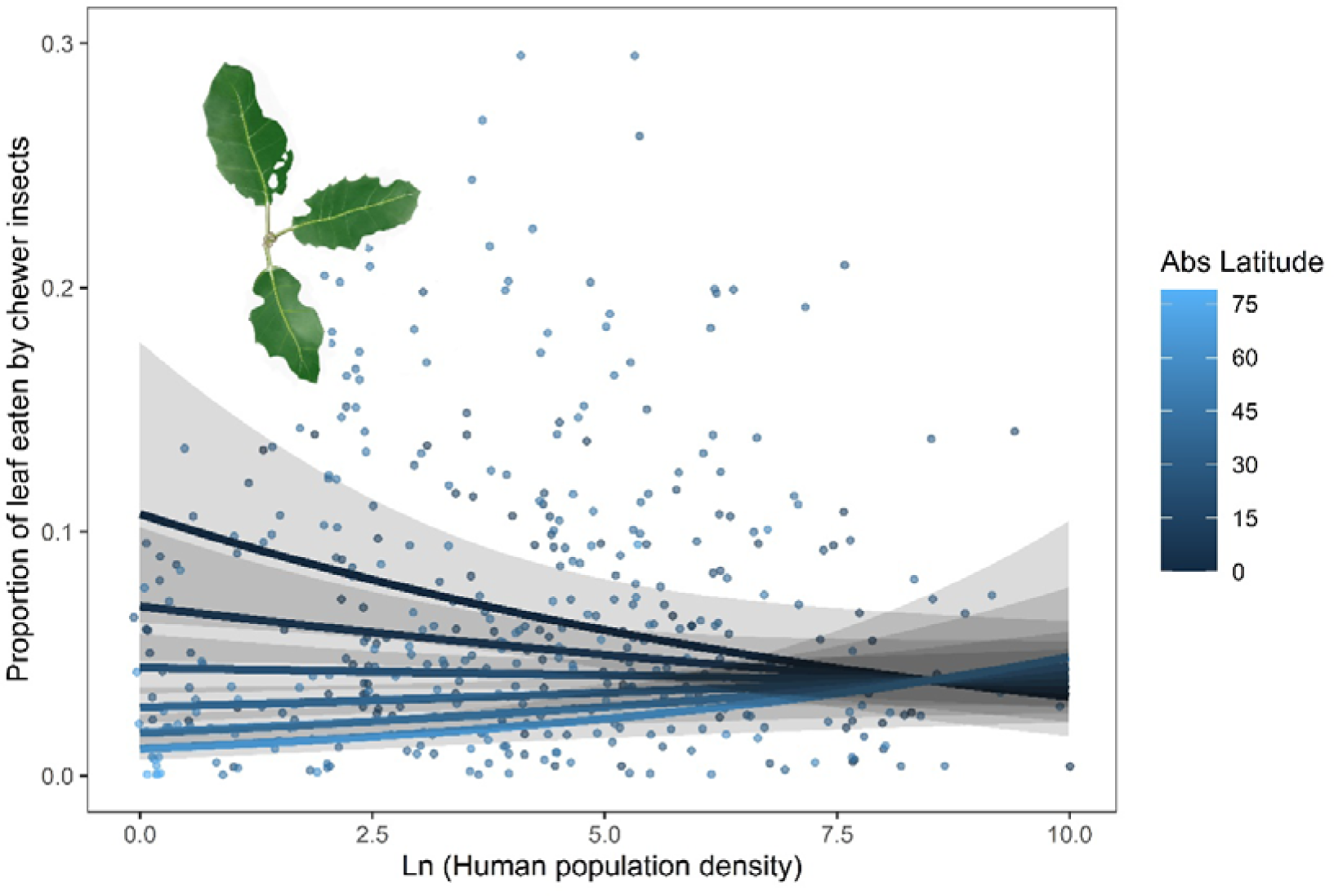
Predictions of generalized linear mixed models showing the proportion of herbivory with 95% confidence intervals along a gradient of human population density (on ln-scale) at six different latitudes (0°, 15°, 30°, 45°, 60° and 75°; marked by differ-ent colors). Mean observed values of the proportion of herbivory per plant species and site are shown by dots. Dots colors represent the absolute latitude.

### Predation

We combined information from 517 different localities spanning 116° latitude (from 42° S to 74° N), with a human population ranging from 0 to 12,401 inhabitants/km^2^. The mean probability of plasticine caterpillar being attacked by a bird during one day of exposure was 0.0447 (CI_95_: 0.0381-0.0512). As with herbivory, there was an effect of human population density on ln-scale, absolute latitude and their interaction on the probability of bird predation (10 km buffer; Table 1; Supplementary material 1 Table S5). We detected spatial autocorrelation in model residuals (Moran I statistic standard deviate = 10.439, p < 0.0001), so best-fit models were re-fitted including a spatial autocorrelation structure.

Model predictions showed changes in the effects of human population on the probability of bird predation along the latitudinal gradient. For lower latitude (< 40°) localities there was a strong decrease on predation as human population increased. This effect weakened towards higher latitudes (subtropical and temperate ecosystems; ∼ 50°) and reversed at higher latitudes (> 60°), i.e. greater predation in more populated areas (Figure 3).

**Figure 3:**
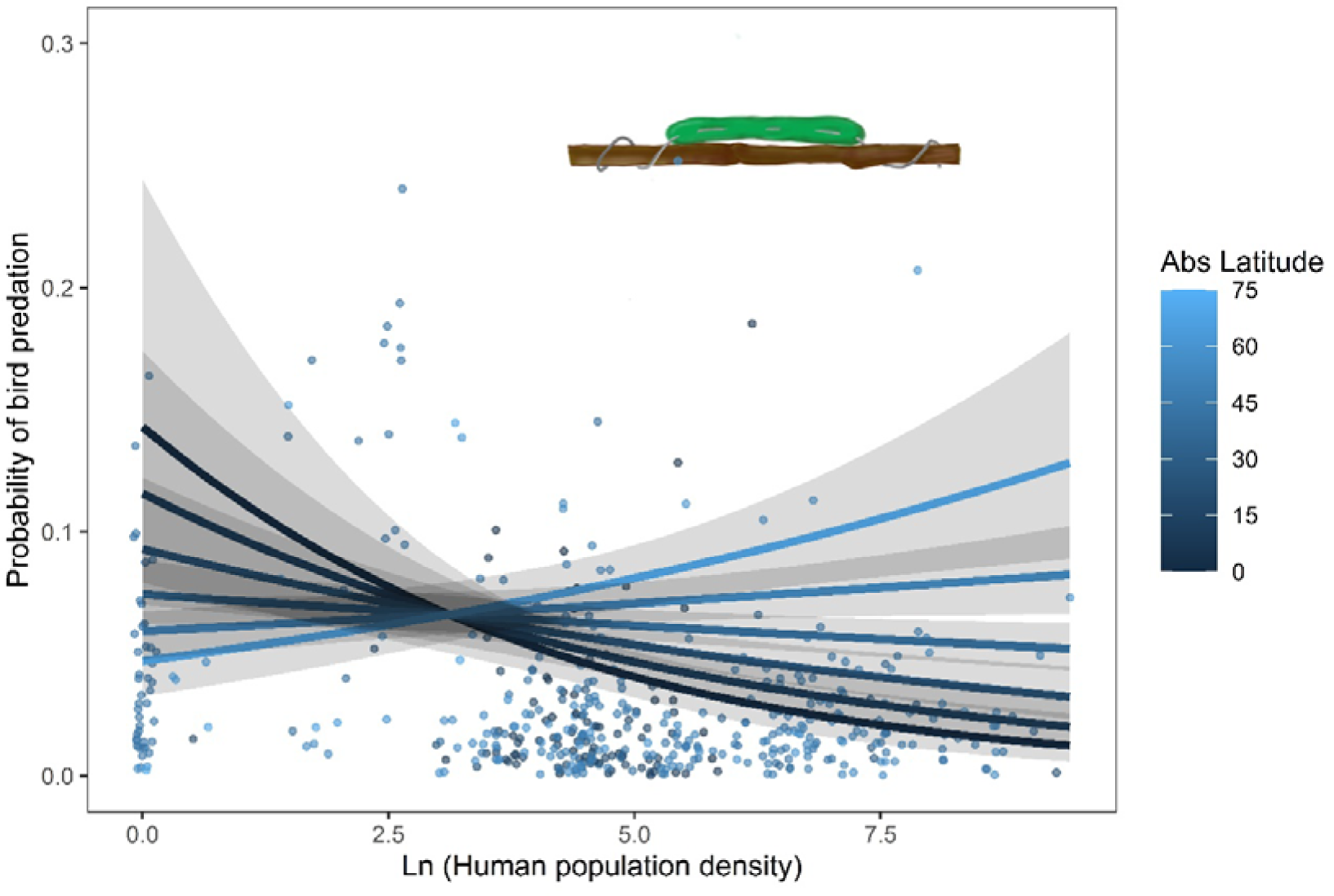
Predictions of generalized linear mixed models showing the probability of bird predation on green caterpillars with 95% confidence intervals along a gradient of human population density (on ln-scale) at six different latitudes (0°, 15°, 30°, 45°, 60° and 75°; marked by different colors). Mean observed values of the probability of bird attack per site are represented as dots. Dots colors represent the absolute latitude.

## Discussion

We expected to find a negative impact of human population on both herbivory and predation, with this effect being less prominent at lower latitudes, where the intensity of biotic interactions is greater than at higher latitudes. Our results partly confirm our initial expectations, but reveal that the urbanization can in fact have a positive effect on the intensity of trophic interactions at higher latitudes. So, whereas in tropical areas we detected the lowest rates of herbivory and predation in highly populated areas, this pattern becomes negligible in temperate regions, and even reverses in polar regions, where herbivory and predation increased with increasing human population density. Thus, the increase of human population density has opposite effects on trophic interaction intensities when comparing high and low latitudes.

The opposite effects of urbanization on herbivory in tropical versus polar latitudes could be explained by the urban heat island effect (Kim, 1992). Cities can be up to 10 °C warmer than surrounding rural areas, and this effect is greater at higher latitudes (Wienert & Kuttler, 2005) and can be easily detected more than 10 km away from the city center (Peng, Hu, Dong, Liu & Liu, 2020). The urban heat island effect might facilitate ectotherm insect activity, particularly at cooler latitudes where minimal temperature limits their performance. Thus, in cities at cooler climates the seasonal period of insect activity can be longer than in sparsely populated areas (Youngsteadt, Dale, Terando, Dunn & Frank, 2015), and this could offset the adverse effects of urbanization found in warmer latitudes (Youngsteadt, Dale, Terando, Dunn & Frank, 2017). The increase of temperature may affect the activity of tropical insects as they tend to be thermal specialists adapted to environmental temperatures, relatively stable throughout the year (Sunday, Bates & Dulvy, 2011). Artificial warmer conditions during winter due to the urban heat island effect could benefit vegetation growth in higher latitudes, while this effect can diminish in lower latitudes where growth is less seasonal (Peng, Ma, Lei, Zhu, Chen et al., 2011). So, whereas at mid latitudes the benefits of increasing temperatures counterbalance the negative influence of human impacts, at higher latitudes the positive effects exceed the negative ones.

The interaction between urbanization and latitude on herbivory has been recently studied both in Europe (Kozlov et al., 2017; Moreira et al., 2019) and the USA (Meineke et al., 2019). Contrary to our results, these studies showed that there was a decrease of chewing herbivory with increasing urban population throughout a latitudinal gradient. The discrepancies between the results reported in these studies and our findings might be related to the larger latitudinal range covered by our study (122°; or 74° of absolute latitude gradient). In addition, some of these studies encompassed smaller ranges in human population density than our study (e.g. Meineke et al., 2019), or included human use categories instead of population density as a surrogate of human impact on the territory (e.g. Kozlov et al., 2017; Moreira et al., 2019), which precludes direct comparison with our results.

The higher abundance of birds and, consequently, the higher levels of bird predation found in cities at high latitudes might be the result of other factors in addition to the urban heat island effect. First, since predator density tends to decrease with increasing latitude (Laurila et al., 2008; Schemske et al., 2009), birds at high latitudes tolerate human presence better than at low latitudes. Second, the stable availability of resources in cities at high latitudes compared to natural areas (Anderies, Katti & Shochat, 2007), results in higher abundances of birds and, consequently, in higher levels of bird predation on insects (Kozlov et al., 2017). And third, because herbivorous insects also benefit from the urban heat island effect, their higher abundance in cities at high latitudes could translate in a higher abundance of insectivorous birds via bottom-up mechanism (Polis, Anderson & Holt, 1997).

## Conclusions

We conclude that changes in trophic interaction intensity with an increase in human population density depend on latitude. Higher human population density at high latitudes increases herbivory and predation pressure, but the opposite trend is observed at low latitudes. Given that cities currently represent temperatures that will be experienced outside of urban areas at the same latitude in the future (Youngsteadt et al., 2015), and based on our results, we could expect: a decrease in the intensity of trophic interactions at tropical latitudes, no effects at temperate latitudes and an increase of trophic interactions in cold latitudes in response to climate change. This could imply important effects on ecosystem services (e.g. nutrient cycling, pest control) that would differently affect tropical and boreal regions, producing a disruption on the natural balance of ecosystems (i.e. ecological imbalance). These differences between cold and tropical latitudes are important since it is predicted that the new big cities (5-10 million people) in the next 10 years will be at tropical latitudes (United Nations, 2018), which will have detrimental consequences for trophic interactions and the ecosystem services they provide. These results open a new venue for studying the effects of human impacts on different ecosystem properties across latitudinal gradients.

## Data Accessibility Statement

All of the data and R scripts used in this study is available at Dryad: “The effects of human population density on trophic interactions are contingent upon latitude” DOI: 10.5061/dryad.sbcc2frcf

