## Supplementary Material 1 for "The effects of human population density on trophic interactions are contingent upon latitude"

Table S1: Data sources on herbivory on woody plants used in this article.

| Source | Range | Date | Sites | Plant species | Plant genera | Plant families |
| --- | --- | --- | --- | --- | --- | --- |
| Kozlov *et al.* (2015) | Global | 2000 - 2014 | 275 | 284 species | 144 genera | 78 families |
| Nakajima (2015) | Japan | 2014 | 22 | 2 species | 1 genus | 1 family |
| Guyot *et al.* (2016) | Europe | 2012 - 2013 | 6 | 15 species | 11 genera | 5 families |
| Valoy et al., (2018) | Argentina | 2018 | 4 | 1 species | 1 genus | 1 family |
| Nakajima (2018) | Japan | 2014 | 9 | 2 species | 2 genus | 1 family |
| Allman *et al.* (2018) | USA | 2012 - 2013 | 1 | 4 species | 1 genus | 1 family |
| Muehleisen et al. (2020) | Panama | 2014 | 3 | 13 species | 13 genus | 11 families |
| Mendes *et al.* (2021) | Global | 2017-2020 | 37 | 193 species | 140 genera | 74 families |
| Wang *et al.* (2022) | China | 2018 | 30 | 1 species | 1 genus | 1 family |
| Hernández-Agüero *et al.* (2023) | Global | 2017-2020 | 48 | 33 species | 22 genera | 20 families |

Table S2: Data sources on bird predation on green plasticine caterpillars used in this article. NP: information was not provided by the authors.

| Source | Range | Date | Sites | Days | Plant species | Attached | Length | Diameter |
| --- | --- | --- | --- | --- | --- | --- | --- | --- |
| Mäntylä *et al.* (2008) | Finland | June 2007 | 1 | 14 | *Betula pubescens* | Metal wire (Ø 0.35 mm) | 2-3 cm | 3-4 mm |
| Mäntylä *et al.* (2014) | Finland | June 2010 | 1 | 14 | *Betula pubescens* | Metal wire (Ø 0.35 mm) | 2-3 cm | 3-4 mm |
| Maas *et al.* (2015) | Indonesia | June to July 2011 | 10 | 4 | *Theobroma cacao* | Power glue | 2 cm | 4 mm |
| Roslin *et al.* (2015) | Global | June 2013 to August 2015 | 31 | 4 | Several species | Power glue | 3 cm | 2.5 mm |
| Kozlov *et al.* (2017) | Europe | May -June 2016 | 4 | 7 | *Betula pubescens* | NP | 3 cm | 4 mm |
| Mrazova & Sam (2018) | Czech Republic | July 2016 | 1 | 4 | *Salix cinerea* | Pinned | 2 cm | 3 mm |
| Roels *et al.* (2018) | Panama | July 2016 | 3 | 10 | Several species | Metal wire | NP | NP |
| Zvereva *et al.* (2019) | Global | May 2017 to December 2018 | 11 | Up to 64 | Several species | Wire (Ø 0.3–0.5 mm) | 1.5-3 cm | 4-5 mm |
| Murray *et al.* (2021) | Malasya | July to August 2016 | 8 | 3 | Several species | Thin wire | NP | NP |
| Schwab *et al.* (2020) | Madagascar | October 2018 | 80 | 2 | Vanilla | Pins | 3.5 cm | 5 mm |
| Zvereva *et al.* (2020) | North Europe | June 2016 to August 2019 | 10 | Up to 17 | *Betula* | Wire (Ø 0.3 mm) | 2.8-3.2 cm | 3.5-4 mm |
| Gomes *et al.* (2021) | USA | May -July 2018 | 48 | 5 | Several species | Power glue | NP | NP |
| Leuenberger *et al.* (2021) | USA | June 2015 | 16 | 6 | *Viburnum lantanoides* and *Fagus grandifolia* | Power glue | 2.2 cm | 3.5 mm |
| Valdés-Correcher *et al.* (2021) | Europe | April 2018 to July 2019 | 255 | 14 | *Quercus robur* | Metal wire (Ø 0.5mm) | 3 cm | 4 mm |
| Zvereva & Kozlov (2021) | Finland | March 2020 | 2 | 7 | *Betula pubescens and Pinus sylvestris* | Wire (Ø 0.3 mm) | 2.5–3 cm | 4–5 mm |
| Hernández-Agüero *et al.* (2023) | Global | May 2017 to August 2020 | 36 | Up to 50 | Several species | Wire (Ø 0.3–0.5 mm) | 1.3-3 cm | 4-5 mm |

Figure S1: Result of the correlation matrix between the human population density calculated for all the study sites of herbivory with radius between 1 to 40 km. There is a representation of the value in a blue-red range.

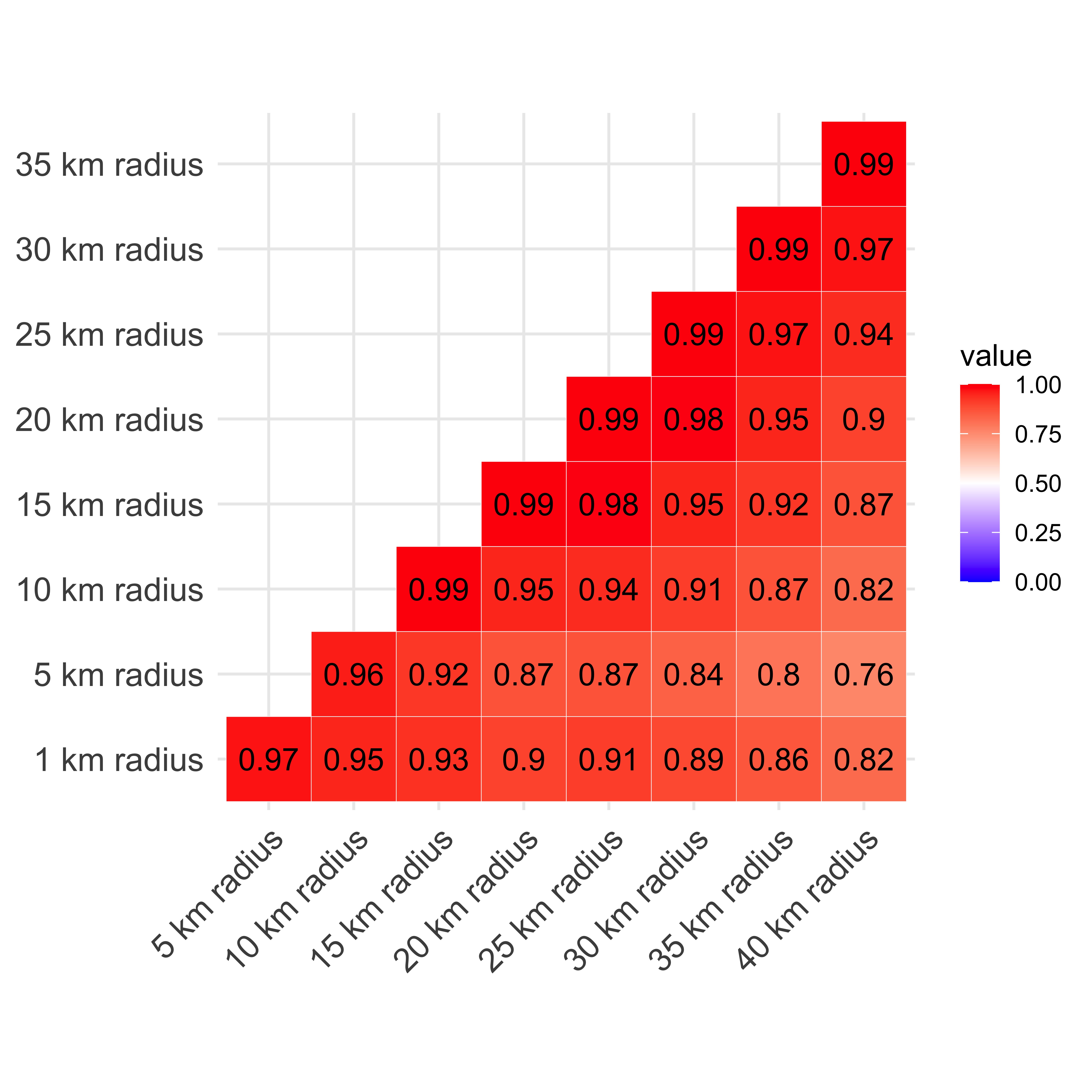

Figure S2: Result of the correlation matrix between the human population density calculated for all the study sites of predation with buffer between 1 to 40 km. There is a representation of the value in a blue-red range.

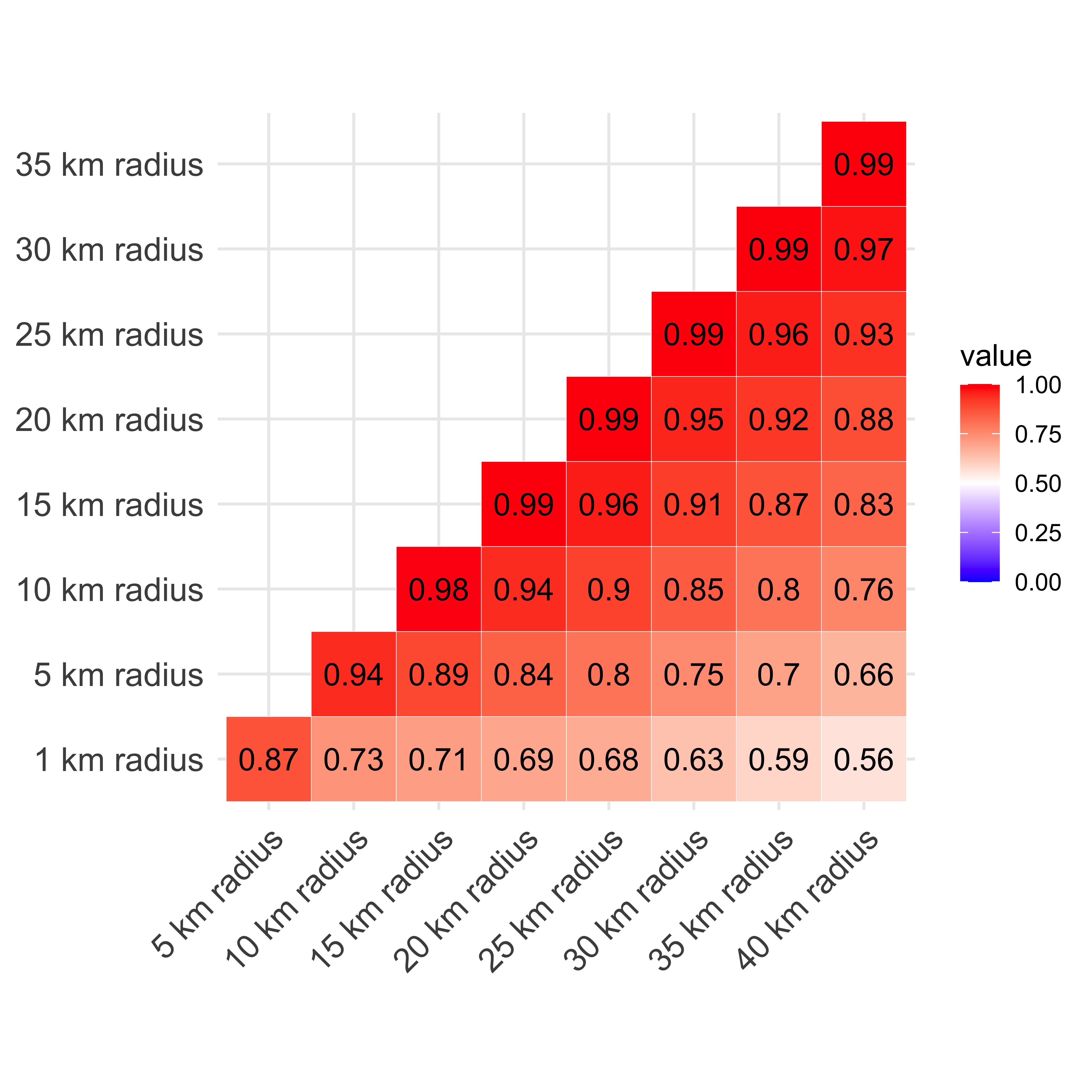

Figure S3: Result of the correlation matrix between latitude, mean temperature, human population density and bult area for all study sites of this study.

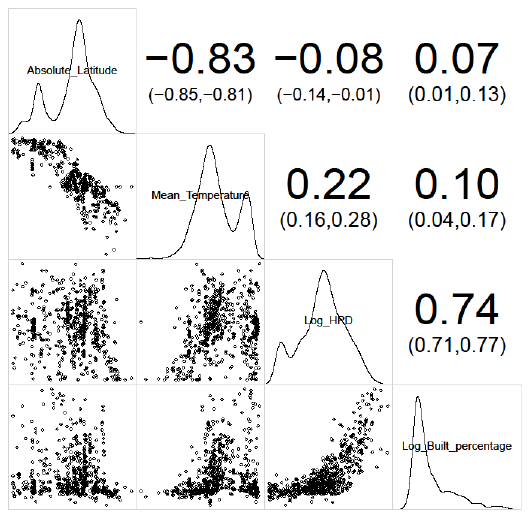

Figure S4: Result of the correlation matrix between latitude, mean temperature, human population density and bult area for herbivory study sites of this study.

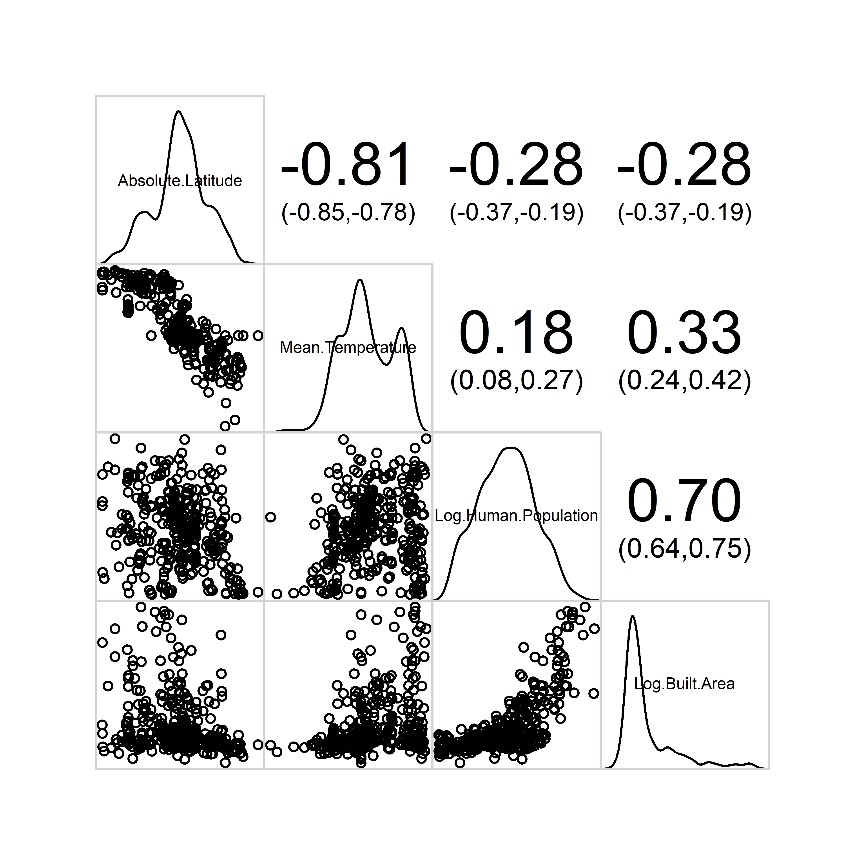

Figure S5: Result of the correlation matrix between latitude, mean temperature, human population density and bult area for predation study sites of this study.

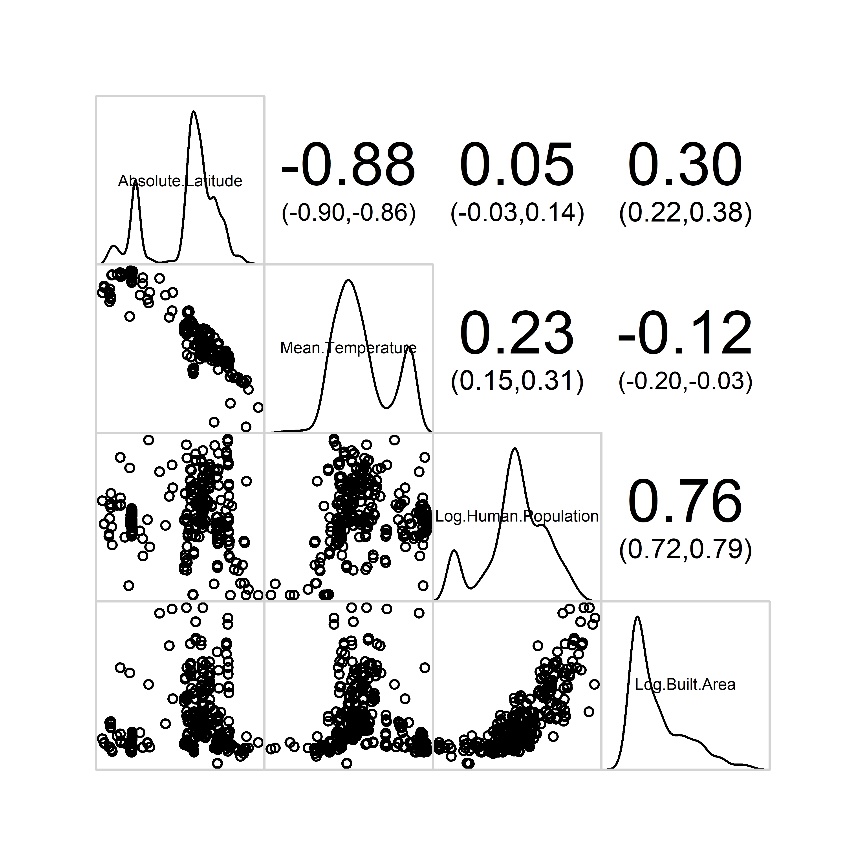

Table S3: The best models for herbivory in different radius indicating the df and the AIC.

| km | Predictors | df | AIC |
| --- | --- | --- | --- |
| 1 | ln(1+Human Population Density) + Absolute Latitude | 7 | -3743.59 |
|  | Absolute Latitude | 6 | -3744.47 |
| 5 | ln(1+Human Population Density) * Absolute Latitude | 8 | -3745.76 |
|  | Absolute Latitude | 6 | -3744.47 |
| 10 | ln(1+Human Population Density) * Absolute Latitude | 8 | -3747.24 |
| 15 | ln(1+Human Population Density) * Absolute Latitude | 8 | -3748.79 |
| 20 | ln(1+Human Population Density) * Absolute Latitude | 8 | -3751.03 |
| 25 | ln(1+Human Population Density) * Absolute Latitude | 8 | -3753.76 |
| 30 | ln(1+Human Population Density) * Absolute Latitude | 8 | -3756.38 |
| 35 | ln(1+Human Population Density) * Absolute Latitude | 8 | -3758.35 |
| 40 | ln(1+Human Population Density) * Absolute Latitude | 8 | -3760.12 |

Table S4: The best models for predation (without including spatial autocorrelation structure) in all the different radiuses indicating the df and the AIC.

| km | Predictors | df | AIC |
| --- | --- | --- | --- |
| 1 | ln(1+Human Population Density) * Absolute Latitude | 6 | -27097.31 |
| 5 | ln(1+Human Population Density) * Absolute Latitude | 6 | -27097.43 |
| 10 | ln(1+Human Population Density) * Absolute Latitude | 6 | -27096.78 |
| 15 | ln(1+Human Population Density) * Absolute Latitude | 6 | -27096.22 |
| 20 | ln(1+Human Population Density) * Absolute Latitude | 6 | -27094.86 |
| 25 | ln(1+Human Population Density) * Absolute Latitude | 6 | -27092.87 |
| 30 | ln(1+Human Population Density) * Absolute Latitude | 6 | -27091.77 |
| 35 | ln(1+Human Population Density) * Absolute Latitude | 6 | -27091.44 |
| 40 | ln(1+Human Population Density) * Absolute Latitude | 6 | -27090.81 |

Table S5: Coefficients of the best models and their standard deviation for both prey-predator and plant-herbivore trophic relations.

|  |  | **Plant-Herbivore** | | **Prey-Predator** | |
| --- | --- | --- | --- | --- | --- |
|  | **Predictor** | **Estimate** | **Std. Dev.** | **Estimate** | **Std. Dev.** |
| **Random** | Site | 0.74772 | 0.8647 | 0.06288 | 0.2508 |
|  | Species/Genus | 0.20918 | 0.4574 |  |  |
|  | Genus | 0.09868 | 0.3141 |  |  |
| **Fixed** | (Intercept) | -2.11953 | 0.299356 | -1.79180 | 0.338243 |
|  | Ln (Human Population) | -0.12771 | 0.057336 | -0.27501 | 0.071893 |
|  | Abs (Latitude) | -0.03172 | 0.006744 | -0.01623 | 0.006621 |
|  | Ln (Human Population) * Abs (Latitude) | 0.003734 | 0.001366 | 0.00521 | 0.001410 |
